# Prioritize diversity or declining species? Trade-offs and synergies in spatial planning for the conservation of migratory birds in the face of land cover change

**DOI:** 10.1101/429019

**Authors:** S Wilson, R. Schuster, A.D. Rodewald, J.R. Bennett, A.C Smith, F.A. La Sorte, P.H. Verburg, P. Arcese

## Abstract

Stemming biodiversity loss requires strategic conservation guided by well-articulated and achievable targets, whether they be proactive (e.g., protect diverse places) or reactive (e.g., protect threatened species). Both types of targets can be effective, but there are trade-offs, especially for broadly-distributed ecosystems or taxa, such as migratory species, a group for which conservation has been challenged by limited knowledge of distributions throughout the annual cycle. We combined novel spatiotemporal distribution models with population trend data to first examine focal areas for the conservation of Neotropical migratory birds (n=112 species) during the non-breeding period in the Western Hemisphere based on a proactive approach (highest diversity) versus a reactive approach (strongest declines) to conservation. For the focal areas, we then assessed the extent of recent anthropogenic impact, protected area status and projected future changes in land cover using three shared socioeconomic pathways (Sustainability=SSP1, Business-as-usual=SSP2, Regional nationalism=SSP3). Spatial priorities were strikingly different when targeting areas of high species diversity, emphasizing southern Mexico and northern Central America, versus areas with more severe declines across species, emphasizing the Andean cordilleras of South America. Only a fraction of the non-breeding region (1.4%) met targets for diversity and decline, mostly in southern Central America. Current levels of protection were similar for the two targets. Areas prioritized to conserve high species diversity have experienced less recent anthropogenic impact than areas prioritized for decline but are predicted to experience more rapid land conversion to less suitable open, agricultural landscapes in the next three decades under both an SSP1 and SSP2 scenario. Only the SSP3 scenario projected similar conversion rates for the two targets. Our findings indicate how even within taxa, efficient conservation efforts will depend on the careful consideration of desired targets combined with reliable predictions about the locations and types of land cover change under alternative socioeconomic futures.

## 1. Introduction

Stemming the unprecedented rates of current biodiversity loss (Ceballos et al. 2017; Pimm et al., 2014) requires strategic and sustained investment in conservation, but socio-economic constraints make difficult choices inevitable (Martin et al., 2018; Wilson et al., 2006). Although sophisticated decision-support tools can inform such choices, the selection of a conservation target is a critical initial step that will profoundly shape outcomes (e.g. Grenyer et al., 2006; Klein et al., 2009; Orme et al., 2005). Potential targets include areas of high species diversity (Buchanan et al., 2011; Somveille et al., 2013), critical habitat for threatened species (ESA, 1973; SARA, 2002), endemism (Myers et al., 2000), or areas of intact wilderness (Watson et al., 2018). Targets may also reflect contrasting conservation paradigms. For example, prioritizing high species diversity may be considered a proactive approach to conserving areas of high biodiversity value, whereas targeting areas with a smaller number of species in rapid decline could be viewed as a more reactive response to species in need of immediate conservation attention (*cf*. Myers et al., 2000; Martin et al., 2018). In this paper, we compare the consequences of proactive vs. reactive approaches for area-based conservation strategies for Neotropical migratory birds during the non-breeding season in the Western Hemisphere.

Migratory species are declining globally (Wilcove and Wikelski, 2008) and present unique challenges in conservation planning (Runge et al., 2014). Because migrants traverse vast distances throughout the annual cycle, we often lack information about which locations and periods of the year most limit population persistence (Rushing et al., 2016; Taylor and Stutchbury, 2016; Wilson et al., 2010; Zurell et al., 2018). Although this knowledge gap creates uncertainty, conservation efforts based upon the best available science must advance as delays in action can imperil populations (Martin et al., 2012). A proactive strategy could still emphasize protection of diverse regions to guard against future threats. In contrast, if the recovery of threatened species is most urgent, then focusing on regions showing the strongest average declines might be most successful in mitigating threats facing species and ecosystems. Of course, focal areas for proactive and reactive targets are not necessarily mutually exclusive, as some areas may have both high diversity and large numbers of declining species (e.g. Hof et al., 2011).

Neotropical migratory birds are the focus of substantial international efforts due to conservation concern for many species (e.g. NABCI, 2016). Most conservation investments for this group have historically funded activities related to the breeding period of the annual cycle in North America and emphasized reactive interventions directed at single species undergoing the strongest declines (ESA, 1973; SARA, 2002). This breeding ground bias has not been entirely deliberate, however, and reflects in part the paucity of information on the non-breeding distribution and ecology of migrants in Neotropical regions. Yet, there is growing recognition and deep concern that rapid land-use change on the non-breeding grounds is a key driver of declines for many threatened Neotropical migrants (Kramer et al., 2018; Taylor and Stutchbury, 2016; Wilson et al., 2018). Advances in crowd-sourced data (e.g. eBird) and species distribution models now allow us to generate weekly estimates of the distribution and abundance for a large number of migratory species over their entire annual cycles (Fink et al., 2010, 2018; La Sorte et al., 2017; Sullivan et al., 2014). By integrating these innovative spatiotemporal distribution models with publicly-available population trend data (e.g., Breeding Bird Survey, Sauer et al., 2016), global databases related to protected areas (WDPA, UNEP-WCMC, 2018) and global land cover change (e.g. Di Minin et al., 2016; Jetz et al., 2007; van Asselen and Verburg, 2013), we can compare proactive and reactive approaches to prioritization and planning and identify potential threats. This comparison will help elucidate trade-offs among approaches that might ultimately affect conservation outcomes under alternative land-use scenarios (Nicholson et al. 2019).

Here, we used weekly estimates of relative abundance for 112 Neotropical migratory birds to accomplish the following objectives:

1. Evaluate differences in the geographic regions and ecosystems targeted for conservation based upon a proactive approach favoring areas of high species diversity vs. a reactive approach emphasizing areas with the strongest average declines across species. This objective included the identification of areas of congruence for the two targets that might allow conservation efforts to enhance both the protection of diversity hotspots and species in need of conservation attention.
2. Compare protected area status, as well as the magnitude and trends in the human footprint (Venter et al., 2016) between focal areas selected for proactive versus reactive conservation targets.
3. Assess projected threats from land-use change within the focal areas using forecasts from a global land systems change model (CLUMondo, van Asselen and Verburg, 2013). To do so, we considered three land-use change scenarios related to regional demands for resources, socioeconomic uncertainties and challenges to mitigation and adaptation (Wolff et al., 2018). Using these land use simulations we examined where current focal areas for proactive and reactive conservation efforts are projected to become less suitable due to land use and climate over the coming decades.

## 2. Materials and Methods

### 2.1. Species selection and study area

We used the eBird citizen-science database (Sullivan et al., 2014) for this analysis. A total of 224 species were available and we identified a subset of these for analysis using the following procedure. We first examined annual eBird distribution maps for all 224 species to identify Neotropical migratory species (n =181 species), defined as those with breeding ranges in North America and non-breeding ranges that extend south of the Tropic of Cancer (Hagan and Johnston, 1992). We then selected terrestrial passerine species that primarily occur in forested or shrubby habitats during the non-breeding season (n =117 species, see Table S1). From these, an additional five species were removed because of insufficient data to accurately model the non-breeding distribution (3 species) or an inability to differentiate congeneric species visually during the non-breeding period (2 species, Table S1). The remaining 112 species sorted into two groups based on the geographic location of their breeding and non-breeding distributions: (1) long-distance migratory species with geographically distinct breeding and non-breeding grounds (*n* = 95), and (2) comparably short-distance migrants with overlapping breeding and non-breeding grounds (*n* = 17).

### 2.2. Estimating species’ distributions and relative abundance

We estimated weekly relative abundance at an 8.34 × 8.34 km spatial resolution for each of the 112 species using spatiotemporal exploratory models (STEM) (Fink et al., 2010, 2018). eBird data used in the STEM include complete checklists collected under the “traveling”, “stationary”, and “areal” protocols from 1 January 2004 to 31 December 2016 within the spatial extent bounded by 180° to 30° W Longitude (as well as Alaska between 150° E and 180° E). This resulted in a dataset consisting of 14 million checklists collected at 1.7 million unique locations. STEM are an ensemble of local regression models generated from a spatiotemporal block subsampling design. Estimates of relative abundance for each species and week are assumed to be stationary. Zero-inflated boosted regression trees are used to predict the observed counts (abundance) of species based on spatial covariates (land cover categories), temporal covariates to account for trends, and predictors that describe the observation/detection process to incorporate variation in detectability. The quantity estimated per 8.34 × 8.34 km pixel was the expected number of birds of a given species by a typical eBird participant on a search starting from the center of the pixel from 7:00 to 8:00 AM while traveling 1 km. For each of the 112 species, we averaged the relative abundance estimates across the weeks during the stationary non-breeding season (defined as 14 November to 14 March) for each of the 8.34 × 8.34 km pixels (Fig. S1). We defined the non-breeding grounds for our analysis as all the 8.34 × 8.34 km pixels (see below) containing five or more of 112 species during the period from 14 November to 14 March. This restriction to five or more species allowed us to include the distributions from all species while excluding large, peripheral areas that are well outside of the main non-breeding range for the vast majority of Neotropical migrants.

### 2.3. Identifying target focal areas

We used STEM estimates of relative abundance for the 112 species to define two conservation targets. Our proactive target was based on species diversity estimated with the Shannon index (Shannon, 1948) for each 8.34 × 8.34 km pixel using species’ estimates of relative abundance in the calculations. We chose the Shannon index as a measure of diversity that included abundance and species richness; across all pixels, estimates of diversity and richness were highly correlated (r = 0.82). Our reactive target was based on the average changes in population size during the breeding season using data from the 1966-2015 North American Breeding Bird Survey (BBS, Environment Canada, 2017; Sauer et al., 2017). Long-term estimates of population trend for Neotropical migratory birds do not exist for the non-breeding period but the continent-wide trends for each species from the breeding period allowed us to estimate the median population trend across all species detected (relative abundance > 0) in each 8.34 × 8.34 km pixel during the non-breeding period. Population trends were unavailable for three species (see Table S1). For each conservation target, we defined focal areas as the top 20% (i.e., based on highest diversity and most negative average population trend). This approach of selecting a top percentage of values in each target category is similar to that used elsewhere (e.g. Hof et al., 2011; Orme et al., 2005). We also identified the geographic regions where the focal areas of the two targets overlapped. To quantify the spatial overlap among our solutions for each target, we used a Bray-Curtis measure of dissimilarity between the focal area 8.34 × 8.34 km pixels for each solution. Dissimilarity calculations were performed using the Vegan package (Oksanen et al., 2015) in R version 3.4.4 (R Core Team, 2018).

### 2.4. Human Footprint

We used the global human footprint index (Venter et al., 2016) to identify recent trends in human pressures for the focal areas selected for each conservation target. The index is a composite measure of human impact derived from eight separate measures: 1) built environments, 2) crop land, 3) pasture land, 4) human population density, 5) night-time lights, 6) railways, 7) roads, and 8) navigable waterways. These eight measures are individually weighted based on their relative levels of human pressure and summed to create a single standardized estimate. The index varies from 0 (no footprint) to 50 (very high footprint) and is estimated at a 1-km spatial resolution across all global terrestrial lands except Antarctica. The index was first measured in 1993 and again in 2009. For all STEM pixels in our analysis, we obtained an average footprint during these two years as well as the change over the 16-yr period. There were approximately 70 footprint estimates for each STEM pixel, which we averaged to create a single measure per pixel.

### 2.5. Protected area coverage

We used the World Database on Protected Areas (WDPA, UNEP-WCMC, 2018) to estimate the extent of protected area coverage for the two conservation targets and the areas of overlap. The WDPA includes seven categories: (Ia) strict nature reserve, (Ib) wilderness area, (II) national park, (III) national monument, (IV) habitat/species management, (V) protected landscape/seascape, (VI) managed resource protected area. Following the same approach as La Sorte et al. (2017), we first combined the WDPA layer with the 8.34 × 8.34 km STEM pixels and identified the protected area category that intersected the center of each pixel. We then calculated the proportion of the land area for each target that contained each of the seven protected area categories. The seven categories were further aggregated into three categories representing high (Ia, Ib), medium (II, III) and low (IV, V, VI) protection status (La Sorte et al., 2017).

### 2.6. Projected land-use change

We used a global land systems map for the year 2000 (Eitelberg et al., 2016; van Asselen and Verburg, 2012) and a global land systems change model (CLUMondo) (van Asselen and Verburg, 2013) to examine land-use change in focal areas for the individual targets and areas of overlap. Spatially explicit land-use change models are important tools to analyze potential land-use trajectories for ecological analysis (e.g. Jetz et al., 2007; LaSorte et al., 2017) and provide information to evaluate policy options. The CLUMondo model simulates land-use change at an approximately 9.3 × 9.3 km spatial resolution based on regional demands for goods and resources dependent on factors that promote or constrain land conversion. Changes in land-use are simulated using empirically quantified relations between land systems, biophysical location and socio-economic factors, in combination with dynamic modeling of competition between different land systems. Model outputs are based on a land systems classification representing combinations of land cover, land use intensity and livestock presence. While the land systems classification in the CLUMondo model includes 17 categories, we aggregated these into six categories for further analysis: (1) forest and mosaic forest-grassland, (2) mosaic forest-cropland, (3) peri-urban and villages (hereafter peri-urban), (4) urban, (5) grassland-bare, (6) cropland or mosaic cropland-grassland (Table S3). The majority of the species considered in our analysis are associated with wooded habitats but many use secondary habitat types including mosaic forest-agriculture and peri-urban landscapes. Open cropland, grassland and bare land cover, in contrast, are likely to contain little to no suitable habitat for these species.

We used the CLUMondo model to simulate land system change for three shared socioeconomic pathway (SSP) scenarios up to 2050. In implementing the three SSP scenarios, model settings are according to the SSP narratives (O’Neill et al., 2014) while demand for agricultural commodities and livestock are derived from assessments with the integrated assessment model IMAGE (Stehfest et al., 2014) at the level of world regions. The Sustainability Scenario (SSP1) and the Regional Nationalism scenario (SSP3) represent contrasting low and high challenges to mitigation and adaptation, respectively (Riahi et al., 2017). In SSP1, development strategies shift globally towards sustainability. Investments in education and health accelerate the demographic transition amid economic growth that focuses more broadly on improving human well-being and reducing inequality among and within countries. Consumption is directed towards low material growth and lower resource and energy intensity. In SSP3, countries experience heightened nationalism, competitiveness and security concerns and regional conflicts that drive a policy agenda oriented toward domestic and regional security issues. Countries focus on achieving energy and food security goals within their own regions at the expense of broader-based development. Population growth is high in developing countries and low in industrialized countries. Environmental concerns remain a low international priority resulting in strong environmental degradation in some regions. The intermediate scenario (Business-as-Usual, SSP2) captures moderate challenges to mitigation and adaptation, with historically consistent trends in technological, economic and societal progress. Population growth continues to rise over the next few decades before leveling off mid-century.

To examine land-use change projections for each target, we aligned the 20% focal areas with the land cover categories in 2000 and the projected land-cover categories under the three SSP scenarios. As a general measure of change in land cover suitability for the Neotropical migrants considered in this analysis, we also identified cases where more suitable land covers containing forest and shrub habitats (i.e., forest, mosaic forest-grassland, mosaic forest-cropland, peri-urban) were projected to become open agricultural lands or barren lands without woody structure (grassland-bare, cropland, mosaic cropland-grassland) under the three SSP scenarios. We excluded land classified as “urban” from analysis because the ability of urban areas to provide habitat to migratory birds is highly variable (e.g., tree cover, green space; Lepczyk et al., 2017; Suarez-Rubio et al., 2013) and urban land comprised only a small proportion of land within our focal areas under all SSP scenarios.

## 3. Results

### 3.1. Geographic variation in target focal areas

Prioritizations based on proactive and reactive targets selected geographically distinct regions (Fig. 1). Focal areas for the proactive target of high diversity (herafter ‘diversity’) were concentrated along coastal and southern Mexico, northern Central America and the western Caribbean (Fig. 1). In contrast, focal areas based on the reactive target of severity of decline (hereafter ‘decline’) were primarily located in the northern Andes of South America with smaller areas elsewhere including the west coast of North America, the Sierra Madre Occidental and the Gran Chaco region of South America (Fig. 1). The spatial overlap between focal areas for the two approaches was remarkably small – only 1.4% (1,325 of 96,078 pixels) of non-breeding locations contained the top 20% of values for both targets (Figure 2, Bray-Curtis dissimilarity = 0.931). Overlapping focal areas were concentrated in southern Central America and northern South America, particularly in the Cordillera Central of Costa Rica, the Cordillera de Talamanca of Costa Rica and west Panama, and the Cordillera Occidental of Colombia (Fig. 2).

**Figure 1.**
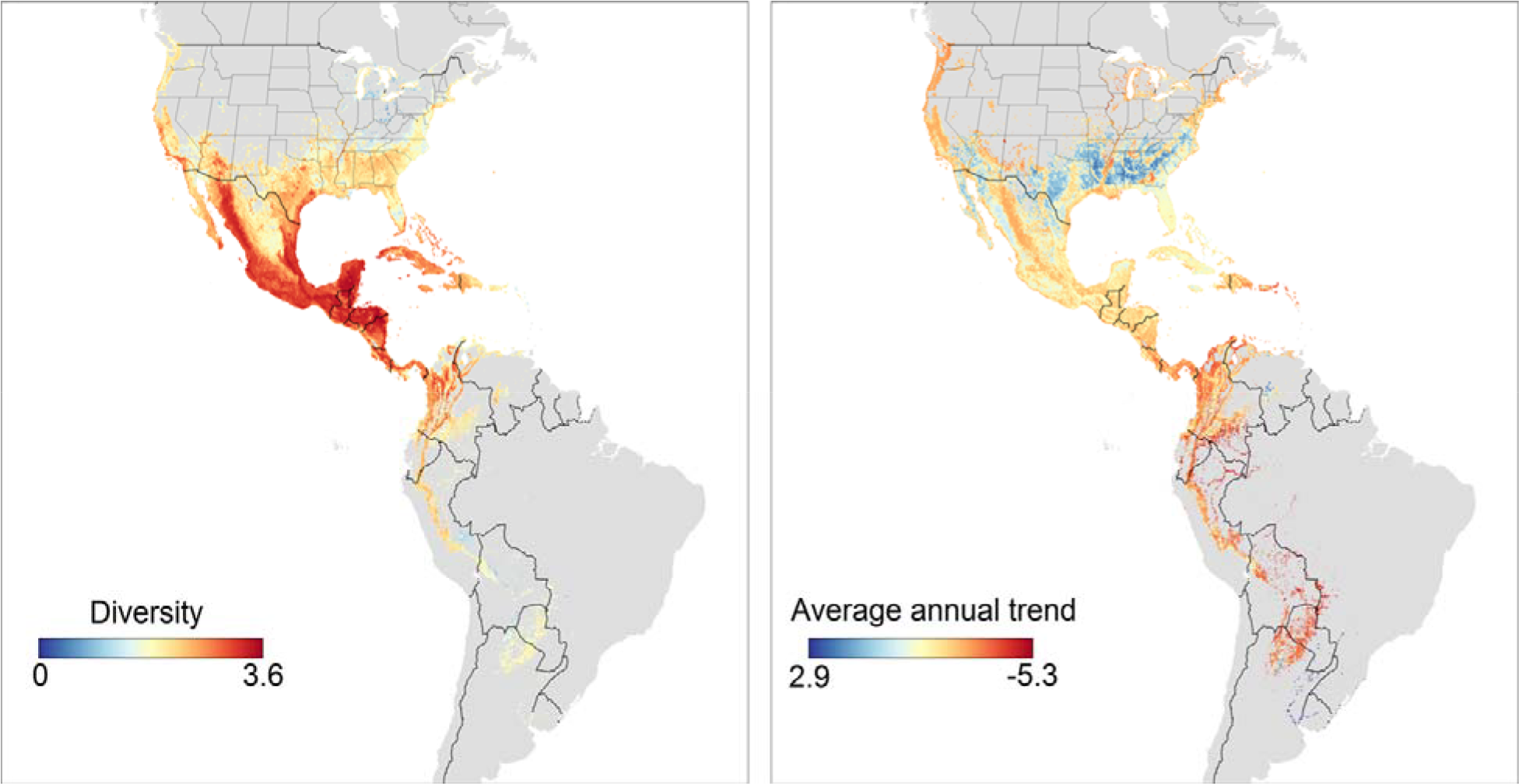
Spatial variation in diversity (left) and average annual trend (right) for 112 Neotropical migratory bird species during the non-breeding season. Diversity is based on the Shannon Index (Shannon, 1948). Average annual trend is the median of the trends between 1966 and 2015 for all species present in each 8.34 × 8.34 km pixel. The non-breeding region was defined by pixels that contained at least 5 species from 14 November to 14 March.

**Figure 2.**
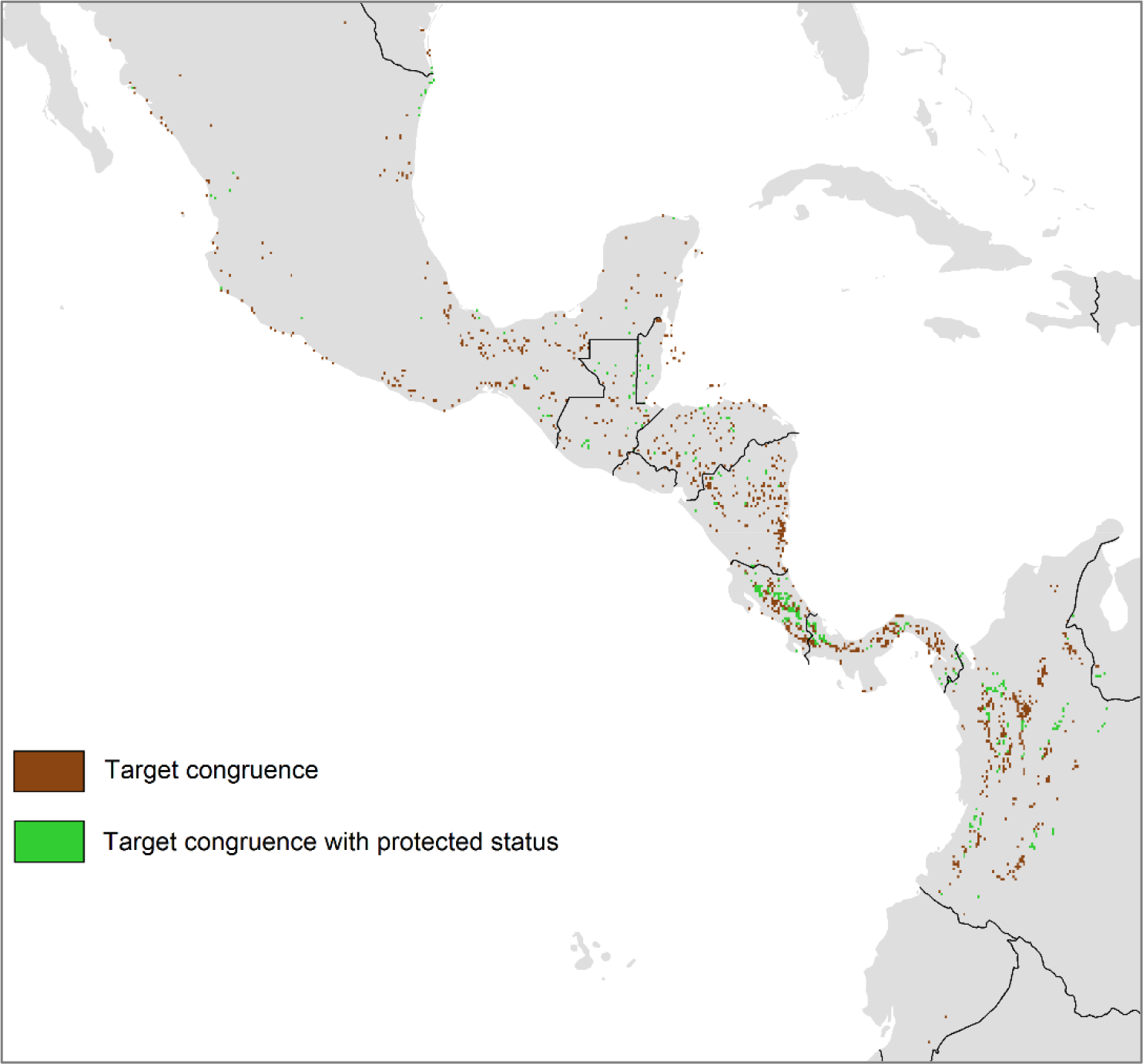
Congruence and current protection of focal areas for high diversity and population decline for 112 Neotropical migratory bird species during the non-breeding season (n = 1,325 overlapping pixels). Green and brown regions are those where the two targets overlap with and without current protected area status, respectively. Focal areas are based on the upper 20% of values for each conservation target.

### 3.2. Recent trends in human footprint among target focal areas

Focal areas for diversity and decline both had a smaller average human footprint than the non-breeding region overall (Table 1). However, they differed in recent trends in footprint; between 1993 and 2009, the index declined by 6.4% in focal areas for diversity but increased by 11.0% in focal areas for decline. This pattern was also reflected in a higher proportion of pixels where the footprint increased between 1993 and 2009 for the decline target (67.2%) than for the diversity target (43.0%). The footprint increased by 16.8% over the same period in areas where the two targets overlapped (Table 1).

**Table 1.** Comparison of trends in the human footprint index for the 20% focal areas selected for each target, areas where the two targets overlap and the non-breeding range used in this analysis (means with 95% CI in brackets). HF = average human footprint index, ∆HF = percent change in HF between 1993 and 2009. The number of selected pixels for each target are shown in brackets after the name.

| Response | HF 1993 | HF 2009 | ΔHF |
| --- | --- | --- | --- |
| Diversity (19,216) | 7.63 (7.53, 7.73) | 7.14 (7.04, 7.25) | -6.38 |
| Severity of decline (19,303) | 5.72 (5.60, 5.85) | 6.35 (6.23, 6.48) | 11.04 |
| Overlap (1,325) | 7.20 (4.87, 9.53) | 8.41 (5.97, 10.85) | 16.79 |
| Region (96,078) | 9.09 (9.04, 9.14) | 9.01 (8.96, 9.06) | -0.88 |

### 3.3. Differences in protected area coverage among focal areas

Across the non-breeding grounds, 15.9% of all pixels had some form of protected status (Table 2). Of the focal areas for diversity, 17.5% were protected areas but with the majority (69.0%) having low protection status, mostly in the form of managed resource areas (Table 2, S1). Focal areas for declines had a similar protected area coverage (16.0%) but in contrast to the diversity target, the majority of these sites (53.0%) had medium protection status and were primarily National Parks (Table 2, S1). Medium status areas in the form of National Parks also represented the majority of protected area classes for areas of congruence for both targets (Fig. 2, Tables 2, S1).

**Table 2.**
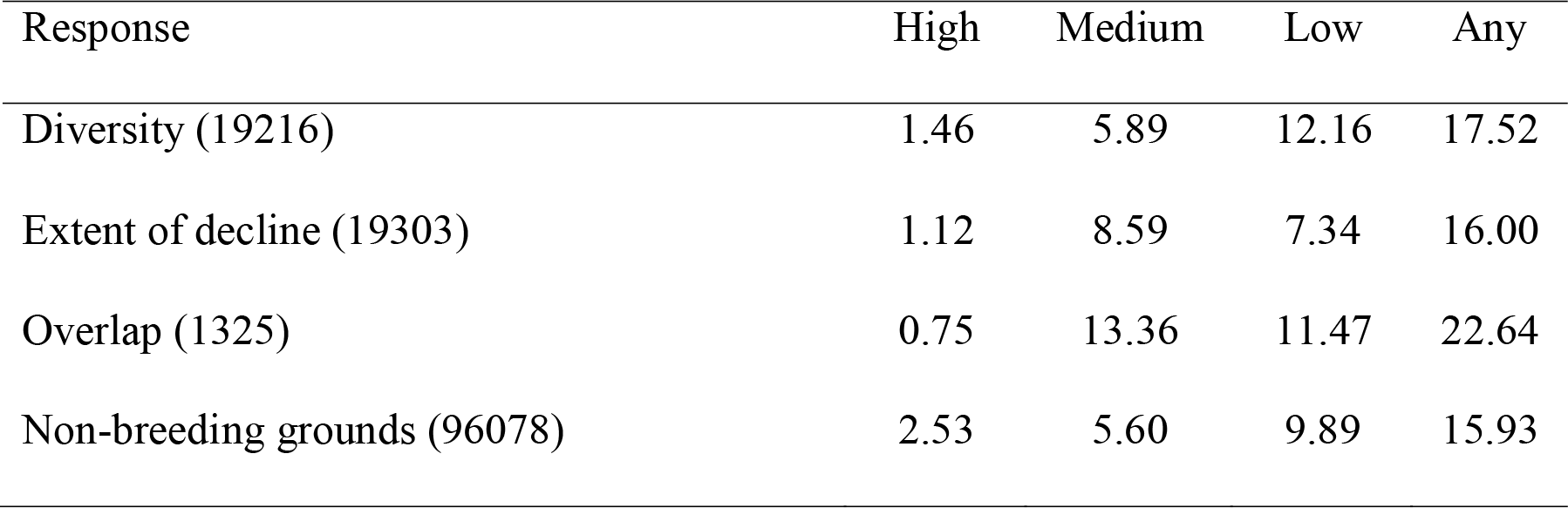
Comparison of protected area coverage for focal areas selected by each target (top 20% of values) based on the World Database on Protected Areas (WDPA, UNEP-WCMC, 2018). Numbers are the percent of 8.34 × 8.34 km pixels for each target that contained high (strict nature reserve, wilderness area), medium (national park, national monument, habitat/species management) and low categories (protected landscape/seascape, managed resource protected area). See Appendix A for the proportion of target pixels for all seven protected area classes.

| Response | High | Medium | Low | Any |
| --- | --- | --- | --- | --- |
| Diversity (19216) | 1.46 | 5.89 | 12.16 | 17.52 |
| Extent of decline (19303) | 1.12 | 8.59 | 7.34 | 16.00 |
| Overlap (1325) | 0.75 | 13.36 | 11.47 | 22.64 |
| Non-breeding grounds (96078) | 2.53 | 5.60 | 9.89 | 15.93 |

### 3.4. Land-cover change

Based on our three SSP scenarios, focal areas that meet proactive diversity targets differed widely in projected land use change (Table 3). Land covers in Pacific coastal regions of Mexico, in particular, transitioned from mixed forest and mosaic forest-cropland to more open and/or intensive land uses – cropland and mosaic cropland-grassland under the sustainability scenario (SSP1) and the regional nationalism scenario (SSP3), and grassland-bare land under the business-as-usual scenario (SSP2) (Fig. 3,Table 3). In contrast, projected changes in land cover were less pronounced for diversity focal areas in the Yucatan and northern Central America, which are expected to remain predominantly forest or mosaic forest under all scenarios (Fig. 3, Figs S2-S5).

**Table 3.** The projected quantity (km^2^× 1000) and percent change from the year 2000 (in brackets) of six land-cover categories in focal areas of high diversity, population declines, and areas of overlap for both targets for 112 Neotropical migratory bird species during the nonbreeding season. Focal areas and overlap are based on the upper 20% of values for each target. The three land-use change scenarios represent the projected quantity of each land-cover category the under low (SSP1), intermediate (SSP2), and high challenge scenarios (SSP3; see Methods for details).

| Target | Scenario | Forest | Mosaic<br>Forest/Crop | Peri-urban /<br>Villages | Urban | Grass/bare | Mosaic<br>Crop/Grass<sup>1</sup> |
| --- | --- | --- | --- | --- | --- | --- | --- |
| Diversity | SSP1 | 657 (-6) | 137 (-51) | 66 (288) | 15 (7) | 48 (-35) | 414 (60) |
| Diversity | SSP2 | 517 (-26) | 39 (-86) | 93 (447) | 15 (7) | 444 (500) | 228 (-12) |
| Diversity | SSP3 | 642 (-8) | 34 (-88) | 98 (477) | 14 (0) | 125 (69) | 424 (64) |
| Decline | SSP1 | 708 (-10) | 62 (-30) | 125 (178) | 32 (28) | 153 (9) | 263 (2) |
| Decline | SSP2 | 486 (-38) | 92 (-5) | 120 (167) | 36 (44) | 309 (119) | 299 (16) |
| Decline | SSP3 | 448 (-43) | 58 (-34) | 170 (278) | 25 (0) | 363 (157) | 279 (9) |
| Overlap | SSP1 | 61 (-6) | 6 (-45) | 8 (300) | 1 (0) | 1 (-50) | 14 (8) |
| Overlap | SSP2 | 49 (-25) | 8 (-27) | 10 (400) | 1 (0) | 12 (500) | 14 (8) |
| Overlap | SSP3 | 49 (-25) | 3 (-73) | 14 (600) | 1 (0) | 9 (350) | 16 (23) |

692 <sup>1</sup> Category includes cropland and mosaic cropland, grassland

**Figure 3.**
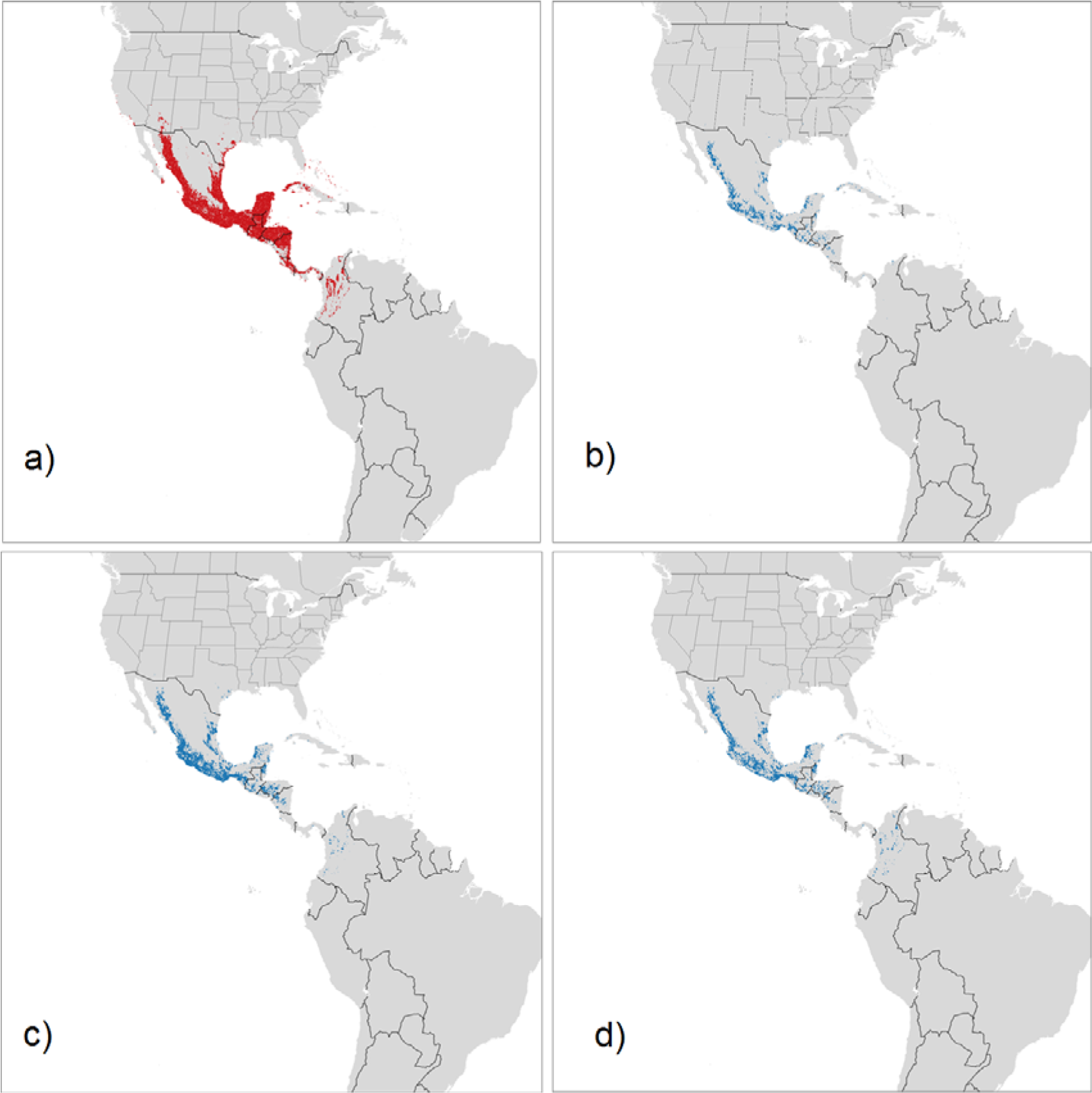
Focal regions for high Neotropical migrant diversity during the non-breeding season (a) and areas within those focal regions where landscapes that were forest, mosaic-forest or peri-urban in 2000 are projected to become open, agricultural landscapes in 2050 under three land use scenarios: Sustainability = SSP1 (b), Business-as-Usual = SSP2 (c), Regional Nationalism = SSP3 (d). Focal regions for high diversity are based on the upper 20% of Shannon index values for 112 Neotropical migratory bird species. See Supplemental Material for figures showing land-cover change under each scenario across the non-breeding region.

The extent and type of projected change in land-cover for focal areas based on average declines also differed by SSP scenario and region (Table 3, Fig. 4). The most extensive differences among scenarios were projected to occur in the cordilleras of northern South America and the Sierra Madre, where land mostly remained in forest or mosaic forest under the SSP1 scenario, but transitioned to open agricultural landscapes under the SSP2 and SSP3 scenarios (Table 3, Fig. 4, Figs S2-S5). Extensive conversion of forest to grassland-bare is also expected under the SSP2 and SSP3 scenarios at the southern limit of species’ non-breeding ranges in the Gran Chaco region (Fig. 4).

**Figure 4.**
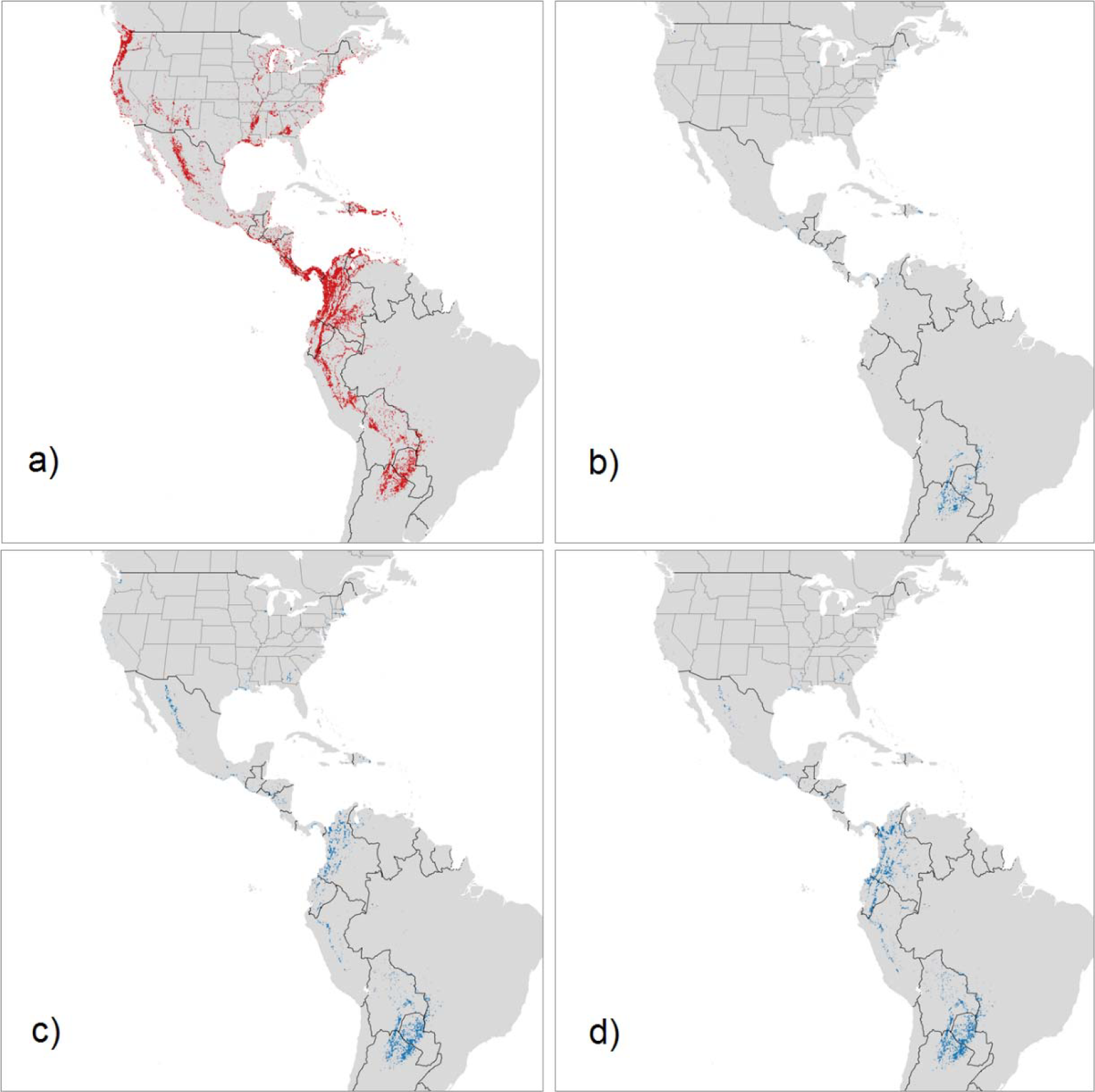
Focal regions for severity of decline for Neotropical migrants during the non-breeding season (a) and areas within those focal regions where landscapes that were forest, mosaic-forest or peri-urban in 2000 are projected to become open, agricultural landscapes in 2050 under three land use scenarios: Sustainability = SSP1 (b), Business-as-Usual = SSP2 (c), Regional Nationalism = SSP3 (d). Focal regions for extent of decline are based on the top 20% of mean, negative population trends for all species in each pixel.

Conversion to less suitable land covers was more substantial for the diversity than decline focal areas (Figs. 3, 4). Of the current focal regions for diversity, conversion from forest, mosaic forest or peri-urban to open agricultural landscapes was predicted for 13% of the region under the SSP1 scenario, 28% under the SSP2 scenario, and 20% under the SSP3 scenario. Conversion percentages for the current focal regions for decline were 7% under the SSP1 scenario, 19% under the SSP2 scenario, and 22% under the SSP3 scenario. Thus, the SSP2 scenario predicted a greater loss of potentially suitable area for focal regions of high diversity while the SSP3 scenario predicted a slightly greater loss for focal regions for decline. For the regions of target overlap, conversion percentages from forest, mosaic-forest or peri-urban to open, agricultural were 5% of the region for the SSP1 scenario, 15% for the SSP2 scenario, and 16% for the SSP3 scenario.

## 4. Discussion

We combined broad-scale estimates of species distribution, abundance, and population trend to identify focal areas for proactive versus reactive conservation approaches for migratory species and by doing so provide a novel tool to predict how land use change may affect conservation strategies and outcomes. Our results emphasize the importance of carefully selecting conservation targets for spatial prioritization as outcomes based on contrasting targets, even for the same species and geographies, may differ profoundly (Klein et al., 2009; Possingham and Wilson 2005). When targeting high species diversity, focal areas were distributed mainly in the northern portion of the non-breeding region in southern Mexico, the Yucatan Peninsula and northern Central America. In contrast, targeting areas with stronger declines emphasized the Andean cordilleras of South America. These findings indicate that even within taxa, efficient conservation planning will depend on clear policy directions on desired targets and reliable predictions about the influence of land cover change on focal species.

The congruence between proactive and reactive conservation targets depends in large part on the degree to which anthropogenic stressors co-occur with diversity hotspots (Hof et al., 2011; Orme et al., 2005; Pimm et al., 2014) as well as the timeframe examined (Bennett and Arcese, 2013). For example, habitat conversion by humans might initially lead to positive spatial correlations between diversity, population decline and human influence, but eventually leads to negative correlations as declining species are extirpated from human-dominated landscapes (Bennett and Arcese, 2013). We observed a slight reduction in the human footprint in focal areas based on high Neotropical migrant diversity, in contrast to the approximately 1% annual increase in footprint in areas targeted for species in greater decline. Even this rapid 1% change in more recent periods understates the long-term extent of habitat loss in the northern Andean region where less than 10% of the original forest cover remains (Henderson et al., 1991). Several recent studies have highlighted the relationship between land conversion in the South American Andes and population declines of Neotropical migrants overwintering in the region (Jones et al., 2004; Kramer et al., 2018, Wilson et al., 2018).

Projections of scenario models for shared socioeconomic pathways (SSP) suggest that focal areas for diversity versus declines differ not only in their past but also likely, future exposure to land conversion. Although focal areas for high diversity experienced less recent change in the human footprint than those for declining species, future land use patterns are expected to change. Under both the sustainable (SSP1) and business-as-usual (SSP2) scenarios, diversity focal areas experienced more land conversion than did decline focal areas. More specifically, 13 and 28% of the forested or partially forested landscapes in diversity focal areas are predicted to be converted to open agricultural or bare landscapes under SSP1 and SSP2 scenarios, in contrast to 7% and 19% of the focal area respectively for the decline target. Only the regional nationalism scenario (SSP3) projected similar conversion rates for the two targets. Projected land conversion within the diversity focal area was largely directed towards one of the most at-risk and intensively used ecosystems in the Neotropics – tropical deciduous and semideciduous forests along the Pacific coast of Mexico (Portillo-Quintero and Sánchez-Azofeifa, 2010; DRYFLOR et al., 2016). This threat is especially concerning given that much of the tropical dry forest in this region is unprotected (Portillo-Quintero and Sánchez-Azofeifa 2010, see also Fig. 2). In contrast to the projected changes along the pacific coast of Mexico, the deciduous forests of the Yucatan and the humid tropical evergreen forests of southern Mexico and the Caribbean slope of northern Central Mexico are expected to remain as forest or mosaic forest over much of the focal area under all three SSP scenarios.

The greater projected land use change towards the western portion of the focal area for diversity highlights the risk of future habitat loss for Neotropical migrants from western North America as many of these species primarily overwinter in western Mexico. Long-term population trends of Neotropical migrants overwintering in this region indicate only slight declines (see Fig. 1), but this risk of habitat loss suggests a potential for future declines and a need to recognize this possibility in current conservation plans. That said, a shift to a sustainable socioeconomic pathway has considerable predicted benefits for both targets with more than a 50% reduction in the conversion of potentially suitable habitats to open, agricultural or bare habitats compared to the business-as-usual or regional nationalism scenarios. This reduction was particularly evident in some regions; for example, among the focal areas for extent of decline, the sustainable pathway scenario (SSP1) retained most of what was projected to be lost under the business-as-usual (SSP2) or regional nationalism (SSP3) scenarios for the Sierra Madre and the Northern Andes although considerable loss was still projected for the Gran Chaco.

Our results carry several caveats. First, we defined focal areas based on the locations representing the upper 20% of values for each target resulting in an area selected of approximately 20-22% of the non-breeding range of all species. This approach of selecting focal areas based on an upper percentile for the target is common (e.g. Grenyer et al., 2006; Orme et al., 2005; Schipper et al., 2008) and the 20% used in our study is similar to the current Convention on Biodiversity efforts to protect 17% of all terrestrial area (SCBD, 2010). However, while the focal areas identified here could be the initial focus we emphasize that higher area protection may be needed for conservation. Moreover, our intention in these efforts is to demonstrate a regional approach identifying potential trade-offs and synergies in identify promising areas for conservation rather than a prescriptive prioritization.

Second, although the Neotropical migratory birds we studied utilize landscapes with woody cover during the non-breeding period, they also vary in their degree of habitat specialization, extent to which they use forests of different age, and sensitivity to landscape-scale loss of forest cover (Petit et al., 1995; Wunderle Jr. and Waide, 1994). While this variation across species may result in a range of responses to land cover change, the conversion of forest and shrub habitat to open crop monocultures (e.g., sun coffee plantations) or pasture would negatively impact nearly all focal species (Céspedes and Bayly, 2018; Donald, 2004; McDermott and Rodewald, 2014). Indeed, the expansion of open, agricultural lands on the wintering grounds is thought to be a principal threat underlying population declines of several Neotropical migrants (Jones et al., 2004; Kramer et al., 2018; Wilson et al., 2018).

Third, because we lack the ability to monitor long-term population trends of Neotropical migrants on the non-breeding grounds, our assessment of regions with a greater extent of decline were based on estimates from breeding bird surveys conducted in North America. This approach matched our intended goal of identifying areas with species in greater decline, but does not account for variation in population trend across the nonbreeding range of the species we considered. In future, emerging methods to estimate population trend in different periods of the annual cycle using eBird data should allow us to refine the spatial analyses reported here.

Finally, our study focused only on the stationary non-breeding period of the annual cycle. While conservation actions could target specific threats during this period, efficient conservation efforts might also consider complementarity across periods of the annual cycle (Zurell et al., 2018). For example, more northern focal areas for high diversity during the stationary non-breeding period might also be priority areas for conservation of South American overwintering species as they pass through northern regions during spring and fall migration.

### Conclusions

Portfolios of sites prioritized using proactive and reactive conservation targets for migratory songbirds differed sharply in terms of geography, ecosystem and habitat types identified, and in their exposure to historic and future threats linked to land use pattern and change. Despite being subject to less anthropogenic land use change historically, areas prioritized to conserve high species diversity are predicted to experience more rapid land conversion in future than areas prioritized to conserve species in decline. These results suggest that proactive approaches have the potential to prevent future declines in the Neotropical migrant birds we studied by helping to keep ‘common species common’ and stemming less severe declines in species currently of low conservation concern but remaining vulnerable (Ceballos et al., 2017; Keith et al., 2015). However, prioritizing areas of high species diversity will largely exclude regions where focal species are currently declining most strongly; many of these species are the focus of species-specific conservation efforts on the breeding grounds (ESA 1973, SARA 2002) but our results point to the importance of protection and restoration of habitat in the northern Andes in particular for the effective conservation of these species. Although our findings represent starting points for decision-making, additional research that includes socio-economic data, projected outcomes of management interventions (e.g., Martin et al., 2018; Naidoo et al., 2006; Wilson et al., 2006), and incorporates resident taxa would improve the ability to identify the most cost-effective and feasible actions for conservation depending on current and projected future land cover changes.

