## Supplemental Figures and Tables for "Prioritize diversity or declining species? Trade-offs and synergies in spatial planning for the conservation of migratory birds in the face of land cover change"

Table S1. Average annual trend for 117 Neotropical migratory passerines considered for analysis. Trends are based on the Breeding Bird Survey calculated between 1966 and 2015 (unavailable for species with a “-”). Species with an asterisk after the common name were left out of the analysis because of insufficient data for the stationary non-breeding range.

| Common name | Scientific name | Trend |
| --- | --- | --- |
| Acadian Flycatcher | *Empidonax virescens* | -0.26 |
| Alder Flycatcher* | *Empidonax alnorum* | -0.89 |
| American Redstart | *Setophaga ruticilla* | -0.28 |
| Ash-throated Flycatcher | *Myiarchus cinerascens* | 1.10 |
| Baltimore Oriole | *Icterus galbula* | -1.49 |
| Bank Swallow | *Riparia riparia* | -5.33 |
| Barn Swallow | *Hirundo rustica* | -1.19 |
| Bay-breasted Warbler | *Setophaga castanea* | -0.25 |
| Bell's Vireo | *Vireo bellii* | 0.63 |
| Black Swift | *Cypseloides niger* | -6.60 |
| Black-and-white Warbler | *Mniotilta varia* | -0.86 |
| Black-billed Cuckoo* | *Coccyzus erythropthalmus* | -1.62 |
| Blackburnian Warbler | *Setophaga fusca* | 0.35 |
| Black-capped Vireo | *Vireo atricapilla* | NA |
| Black-chinned Hummingbird | *Archilochus alexandri* | 1.29 |
| Black-headed Grosbeak | *Pheucticus melanocephalus* | 0.72 |
| Blackpoll Warbler | *Setophaga striata* | -4.85 |
| Black-throated Blue Warbler | *Setophaga caerulescens* | 1.95 |
| Black-throated Green Warbler | *Setophaga virens* | 0.35 |
| Black-throated Grey Warbler | *Setophaga nigrescens* | -1.26 |
| Blue Grosbeak | *Passerina caerulea* | 0.81 |
| Blue-grey Gnatcatcher | *Polioptila caerulea* | 0.38 |
| Blue-headed Vireo | *Vireo solitarius* | 2.86 |
| Blue-winged Warbler | *Vermivora cyanoptera* | -0.70 |
| Broad-tailed Hummingbird | *Selasphorus platycercus* | -1.45 |
| Bullock's Oriole | *Icterus bullockii* | -0.66 |
| Calliope Hummingbird | *Selasphorus calliope* | -0.23 |
| Canada Warbler | *Cardellina canadensis* | -2.05 |
| Cape May Warbler | *Setophaga tigrina* | -2.51 |
| Cassin's Kingbird | *Tyrannus vociferans* | 0.19 |
| Cassin's Vireo | *Vireo cassinii* | 1.08 |
| Cave Swallow | *Petrochelidon fulva* | 21.25 |
| Cerulean Warbler | *Setophaga cerulea* | -2.63 |
| Chestnut-sided Warbler | *Setophaga pensylvanica* | -1.15 |
| Chimney Swift | *Chaetura pelagica* | -2.50 |
| Cliff Swallow | *Petrochelidon pyrrhonota* | 0.72 |
| Common Nighthawk | *Chordeiles minor* | -1.93 |
| Common Yellowthroat | *Geothlypis trichas* | -1.01 |
| Connecticut Warbler* | *Oporornis agilis* | -1.93 |
| Cordilleran Flycatcher | *Empidonax occidentalis* | -0.30 |
| Dusky Flycatcher | *Empidonax oberholseri* | -0.60 |
| Eastern Kingbird | *Tyrannus tyrannus* | -1.28 |
| Eastern Wood Pewee | *Contopus virens* | -1.40 |
| Golden-cheeked Warbler^+^ | *Setophaga chrysoparia* | -1.30 |
| Golden-winged Warbler | *Vermivora chrysoptera* | -2.28 |
| Grace's Warbler | *Setophaga graciae* | -1.50 |
| Great Crested Flycatcher | *Myiarchus crinitus* | -0.03 |
| Grey Catbird | *Dumetella carolinensis* | -0.01 |
| Grey Vireo | *Vireo vicinior* | 2.10 |
| Grey-cheeked Thrush | *Catharus minimus* | -2.65 |
| Hammond's Flycatcher | *Empidonax hammondii* | 0.75 |
| Hepatic Tanager | *Piranga hepatica* | 1.34 |
| Hermit Warbler | *Setophaga occidentalis* | -0.05 |
| Hooded Oriole | *Icterus cucullatus* | 0.51 |
| Hooded Warbler | *Setophaga citrina* | 1.36 |
| Indigo Bunting | *Passerina cyanea* | -0.73 |
| Kentucky Warbler | *Geothlypis formosa* | -0.90 |
| Lazuli Bunting | *Passerina amoena* | 0.21 |
| Least Flycatcher | *Empidonax minimus* | -1.71 |
| Louisiana Waterthrush | *Parkesia motacilla* | 0.60 |
| Lucy's Warbler | *Leiothlypis luciae* | 0.75 |
| MacGillivray's Warbler | *Geothlypis tolmiei* | -1.66 |
| Magnolia Warbler | *Setophaga magnolia* | 0.87 |
| Mourning Warbler | *Geothlypis philadelphia* | -1.18 |
| Nashville Warbler | *Leiothlypis ruficapilla* | 0.01 |
| Northern Parula | *Setophaga americana* | 1.11 |
| Northern Rough-winged Swallow | *Stelgidopteryx serripennis* | -0.53 |
| Northern Waterthrush | *Parkesia noveboracensis* | 1.19 |
| Olive-sided Flycatcher | *Contopus cooperi* | -3.10 |
| Orange-crowned Warbler | *Leiothlypis celata* | -0.61 |
| Orchard Oriole | *Icterus spurius* | -0.87 |
| Ovenbird | *Seiurus aurocapilla* | -0.07 |
| Pacific-slope Flycatcher | *Empidonax difficilis* | -0.30 |
| Painted Bunting | *Passerina ciris* | -0.12 |
| Painted Whitestart | *Myioborus pictus* | NA |
| Palm Warbler | *Setophaga palmarum* | -1.78 |
| Philadelphia Vireo | *Vireo philadelphicus* | 2.31 |
| Plumbeous Vireo | *Vireo plumbeus* | -2.44 |
| Prairie Warbler | *Setophaga discolor* | -1.85 |
| Prothonotary Warbler | *Protonotaria citrea* | -1.10 |
| Purple Martin | *Progne subis* | -0.91 |
| Red-eyed Vireo | *Vireo olivaceus* | 0.75 |
| Red-faced Warbler | *Cardellina rubrifrons* | NA |
| Rose-breasted Grosbeak | *Pheucticus ludovicianus* | -0.86 |
| Ruby-crowned Kinglet | *Regulus calendula* | 0.47 |
| Ruby-throated Hummingbird | *Archilochus colubris* | 1.44 |
| Rufous Hummingbird | *Selasphorus rufus* | -1.98 |
| Scarlet Tanager | *Piranga olivacea* | -0.22 |
| Scissor-tailed Flycatcher | *Tyrannus forficatus* | -0.78 |
| Scott's Oriole | *Icterus parisorum* | -0.83 |
| Summer Tanager | *Piranga rubra* | 0.22 |
| Swainson's Thrush | *Catharus ustulatus* | -0.84 |
| Swainson's Warbler | *Limnothlypis swainsonii* | 1.20 |
| Tennessee Warbler | *Leiothlypis peregrina* | -1.03 |
| Townsend's Warbler | *Setophaga townsendi* | -0.60 |
| Tree Swallow | *Tachycineta bicolor* | -1.38 |
| Vaux's Swift | *Chaetura vauxi* | -1.81 |
| Veery* | *Catharus fuscescens* | -1.13 |
| Violet-green Swallow | *Tachycineta thalassina* | -0.66 |
| Virginia's Warbler | *Leiothlypis virginiae* | -1.36 |
| Warbling Vireo | *Vireo gilvus* | 0.85 |
| Western Kingbird | *Tyrannus verticalis* | 0.06 |
| Western Tanager | *Piranga ludoviciana* | 1.28 |
| Western Wood Pewee | *Contopus sordidulus* | -1.37 |
| White-eyed Vireo | *Vireo griseus* | 0.62 |
| White-throated Swift | *Aeronautes saxatalis* | -1.68 |
| Willow Flycatcher* | *Empidonax traillii* | -1.48 |
| Wilson's Warbler | *Cardellina pusilla* | -1.80 |
| Wood Thrush | *Hylocichla mustelina* | -1.91 |
| Worm-eating Warbler | *Helmitheros vermivorum* | 0.38 |
| Yellow Warbler | *Setophaga aestiva* | -0.61 |
| Yellow-bellied Flycatcher | *Empidonax flaviventris* | 2.26 |
| Yellow-billed Cuckoo | *Coccyzus americanus* | -1.45 |
| Yellow-breasted Chat | *Icteria virens* | -0.62 |
| Yellow-rumped Warbler | *Setophaga coronata* | -0.40 |
| Yellow-throated Vireo | *Vireo flavifrons* | 0.98 |
| Yellow-throated Warbler | *Setophaga dominica* | 0.98 |

^+^ based on estimates in The Birds of North America online (Ladd and Gass, 1999)

Ladd, C. and L. Gass (1999). Golden-cheeked Warbler (Setophaga chrysoparia), version 2.0. In The Birds of North America (A. F. Poole and F. B. Gill, Editors). Cornell Lab of Ornithology, Ithaca, NY, USA. <https://doi.org/10.2173/bna.420>

Table S2. Comparison of current protected area coverage for hotspot landscapes selected by each target. Values in the table show the percent of selected landscapes for each target that contained each protected area type: (IA) strict nature reserve, (IB) wilderness area, (II) national park, (III) national monument, (IV) habitat/species management, (V) protected landscape/seascape, (VI) managed resource protected area (WDPA, UNEP-WCMC 2018).

| Response | Ia | Ib | II | III | IV | V | VI | Any |
| --- | --- | --- | --- | --- | --- | --- | --- | --- |
| Diversity (19216) | 1.29 | 0.17 | 4.29 | 0.61 | 0.99 | 0.54 | 11.62 | 17.52 |
| Severity of decline (19303) | 0.32 | 0.81 | 7.15 | 0.42 | 1.02 | 2.45 | 4.89 | 16.00 |
| Overlap (1325) | 0.75 | 0.00 | 10.72 | 1.06 | 1.58 | 0.45 | 11.02 | 22.64 |
| Non-breeding region (96078) | 0.50 | 2.03 | 3.83 | 0.46 | 1.31 | 3.66 | 6.23 | 15.93 |

Table S3.

| **Original Classification** | **New Classification** |
| --- | --- |
| Cropland; extensive with few livestock | Mosaic cropland/grassland or cropland (6) |
| Cropland; extensive with bovines, goats & sheep | Mosaic cropland/grassland or cropland (6) |
| Cropland; medium intensive with few livestock | Mosaic cropland/grassland or cropland (6) |
| Cropland; medium intensive with bovines, goats & sheep | Mosaic cropland/grassland or cropland (6) |
| Cropland; intensive with few livestock | Mosaic cropland/grassland or cropland (6) |
| Cropland; intensive with bovines, goats & sheep | Mosaic cropland/grassland or cropland (6) |
| Mosaic cropland and grassland with bovines, goats & sheep | Mosaic cropland/grassland or cropland (6) |
| Mosaic cropland (extensive) and grassland with few livestock | Mosaic cropland/grassland or cropland (6) |
| Mosaic cropland (medium intensive) and grassland with few livestock | Mosaic cropland/grassland or cropland (6) |
| Mosaic cropland (intensive) and grassland with few livestock | Mosaic cropland/grassland or cropland (6) |
| Mosaic cropland (extensive) and forest with few livestock | Mosaic forest/cropland (2) |
| Mosaic cropland (medium intensive) and forest with few livestock | Mosaic forest/cropland (2) |
| Mosaic cropland (intensive) and forest with few livestock | Mosaic forest/cropland (2) |
| Dense forest | Forest (1) |
| Open forest with few livestock | Forest (1) |
| Mosaic grassland and forest | Forest (1) |
| Mosaic grassland and bare | Grassland/bare (5) |
| Natural grassland | Grassland/bare (5) |
| Grassland with few livestock | Grassland/bare (5) |
| Grassland with bovines, goats and sheep | Grassland/bare (5) |
| Bare | Grassland/bare (5) |
| Bare with few livestock | Grassland/bare (5) |
| Peri-urban & villages | Peri-urban (3) |
| Urban | Urban (4) |

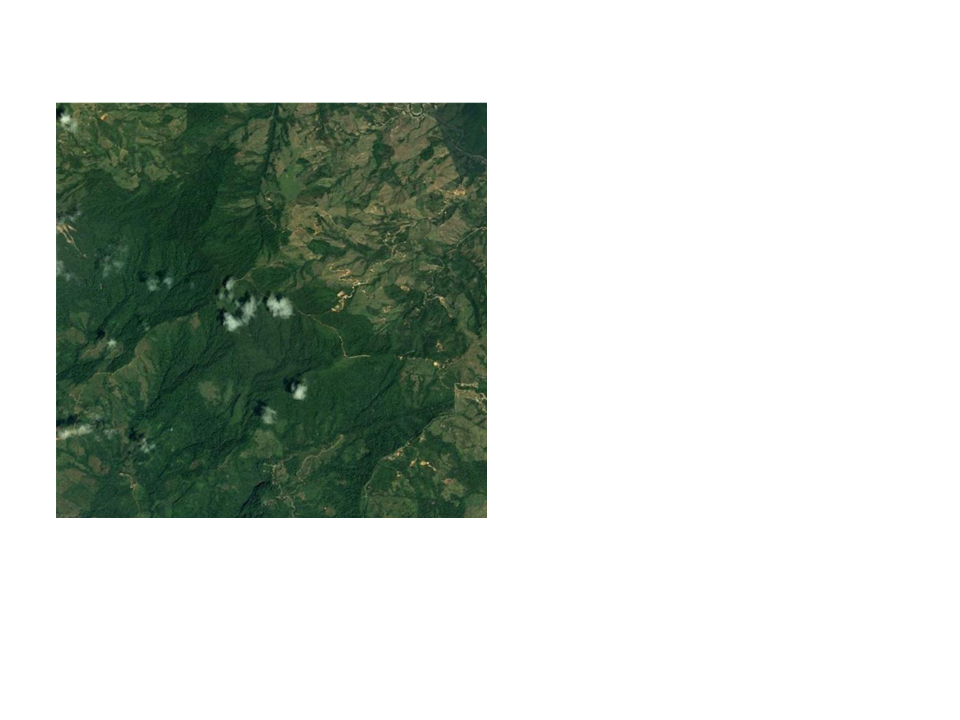

Figure S1. An example eBird pixel approximately 8.3 × 8.3 km in southwest Costa Rica. Using STEM distribution maps for 112 Neotropical migrants we predicted that this landscape would have a Shannon diversity = 3.04 and a median annual population decline across species = 0.85% per year between 1966 and 2015. We estimated these metrics for 96,079 landscapes across the non-breeding range of Neotropical migrants in the western hemisphere.

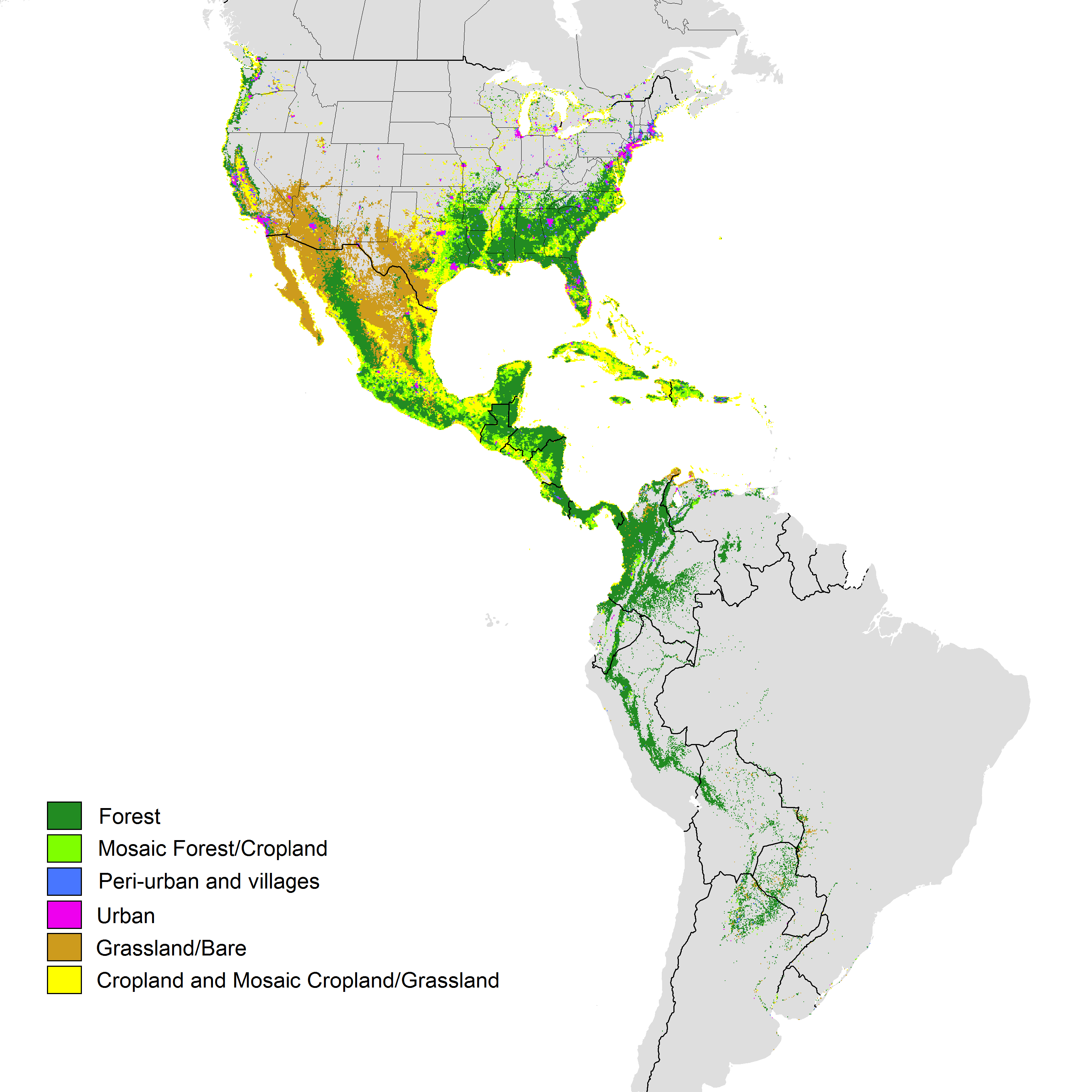

Figure S2. Land cover in 2000 for the stationary non-breeding region of 112 Neotropical migrant birds. All regions selected here contained ≥ 5 of the Neotropical migrant species considered in this analysis.

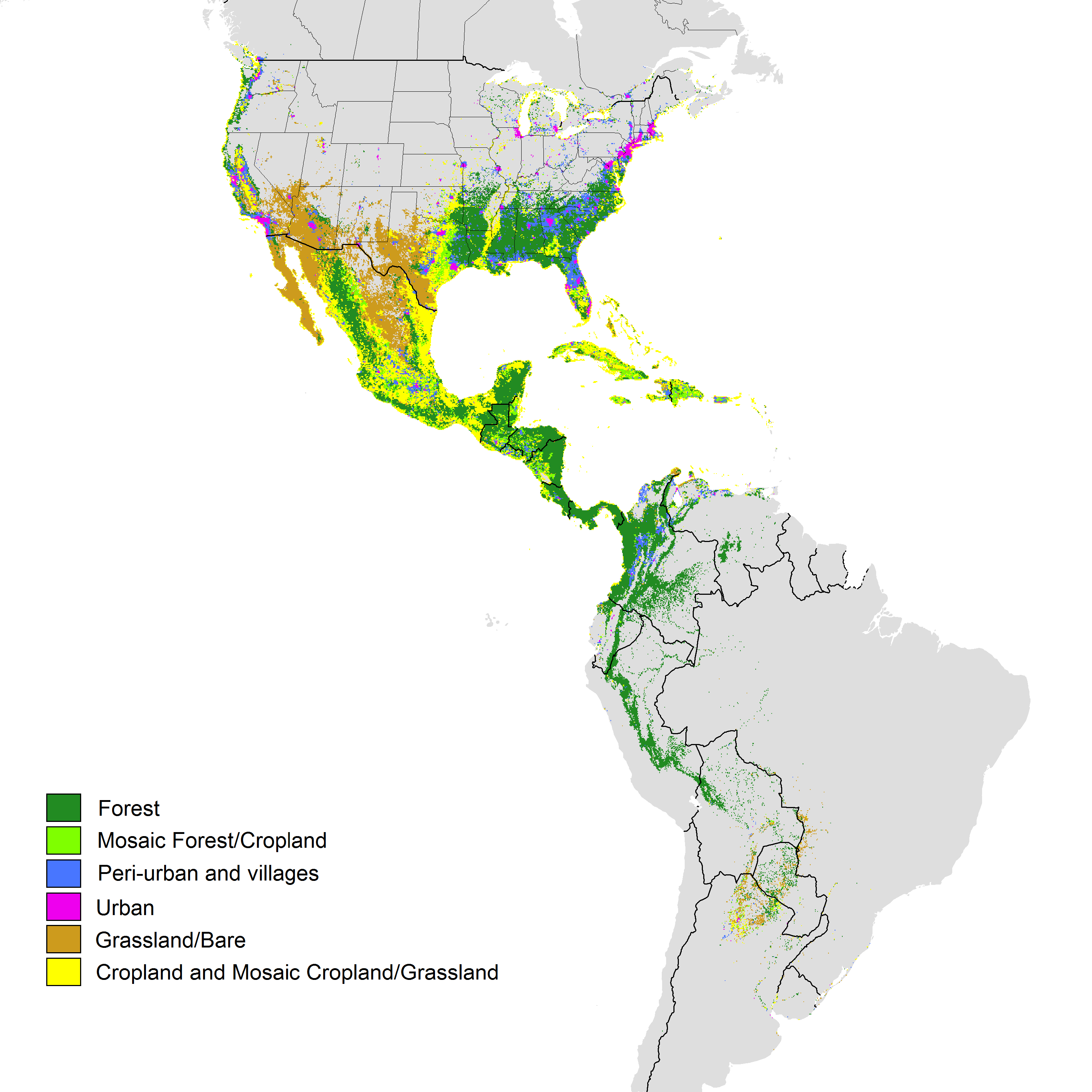

Figure S3. Predicted land cover in 2050 for the stationary non-breeding region of 112 Neotropical migrant birds under shared socio-economic pathway (SSP) 1 emphasizing a sustainability scenario with low challenges to mitigation and adaptation. All regions selected here contained ≥ 5 of the Neotropical migrant species considered in this analysis.

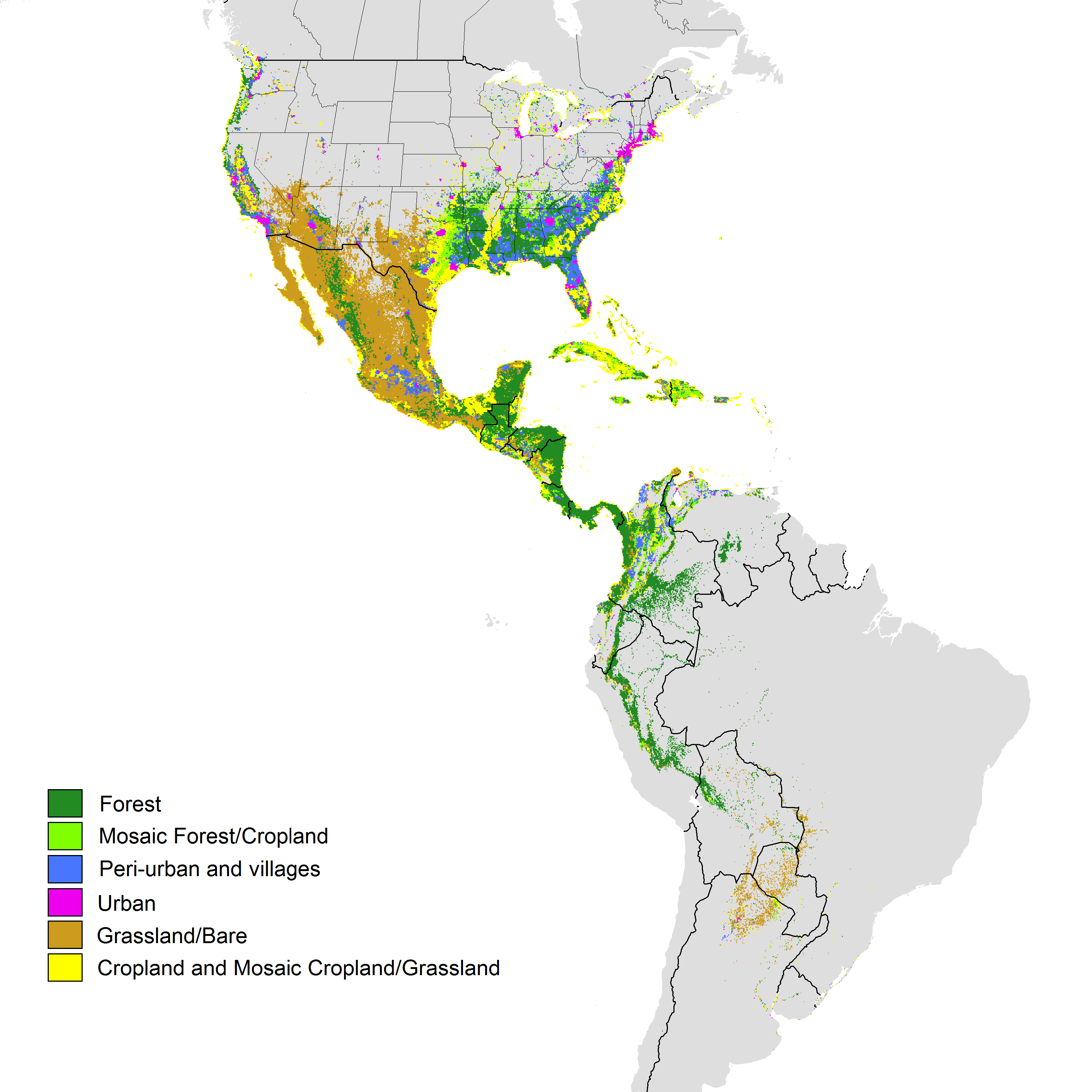

Figure S4. Predicted land cover in 2050 for the stationary non-breeding region of 112 Neotropical migrant birds under shared socio-economic pathway (SSP) 2 emphasizing a business as usual scenario with moderate challenges to mitigation and adaptation. All regions selected here contained ≥ 5 of the Neotropical migrant species considered in this analysis.

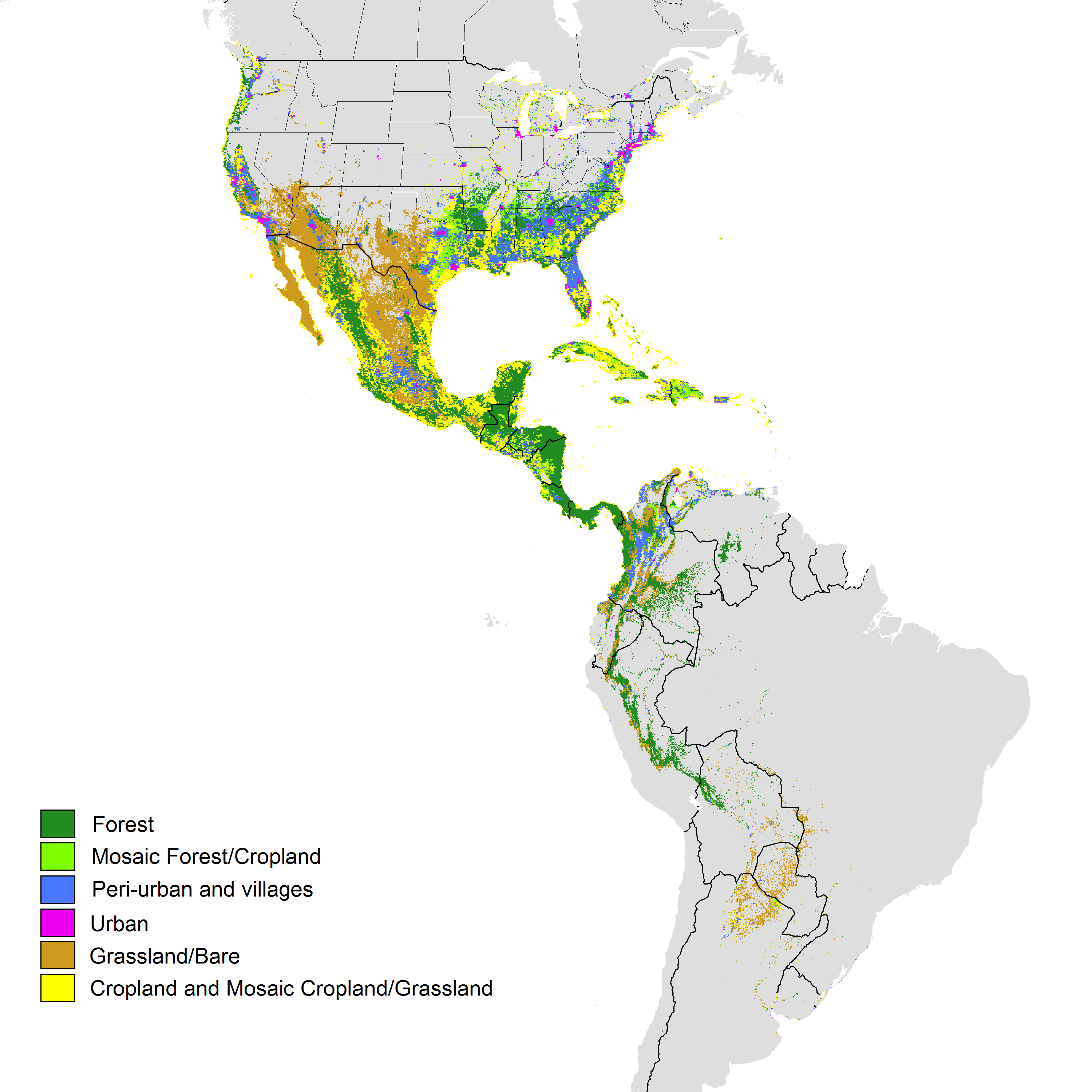

Figure S5. Predicted land cover in 2050 for the stationary non-breeding region of 112 Neotropical migrant birds under shared socio-economic pathway (SSP) 3 emphasizing a regional nationalism scenario with high challenges to mitigation and adaptation. All regions selected here contained ≥ 5 of the Neotropical migrant species considered in this analysis.
